# CellExLink: End-to-end cell-type recognition and normalization in biomedical text

**DOI:** 10.64898/2026.05.26.728013

**Authors:** Alimire Nabijiang, Leili Shahriyari

## Abstract

Since cells are the main components of many biological and biomedical studies, cell-type extraction is an important task in biomedical text mining. However, current biomedical text-mining systems either do not explicitly support cell-type extraction, provide limited support for Cell Ontology normalization, or show limited performance in end-to-end cell-type extraction. These limitations can affect downstream tasks that depend on reliable cell-type information. Here, we present CellExLink, an end-to-end biomedical natural language processing pipeline designed specifically for cell-type recognition and Cell Ontology normalization in biomedical text. The pipeline is designed to improve extraction accuracy and practical usability in literature-mining workflows, while accounting for computational efficiency in its recognition and normalization design. We evaluate CellExLink across heterogeneous biomedical corpora and compare it with established and recent biomedical text-mining tools. The results show that CellExLink provides reliable cell-type recognition, Cell Ontology normalization, and end-to-end extraction across these corpora. By addressing the need for reliable end-to-end cell-type recognition and Cell Ontology normalization, CellExLink can support downstream tasks such as curation, search, relation extraction, and knowledge graph construction.

**Author summary:** Cell types are central to biomedical research, but biomedical papers often use different names, abbreviations, and synonyms for the same cell type. This variation makes it difficult for automated processes to collect and compare cell-type information across papers. Reliable automated extraction is important because literature mining requires consistent cell-type identification before evidence from different studies can be searched, integrated, or reused. Existing off-the-shelf biomedical text-mining tools provide useful functionality, but their ability to support cell-type extraction remains limited and inconsistent. To address this gap, we developed CellExLink, a pipeline that finds cell-type entities in biomedical text and links them to standard Cell Ontology identifiers. We evaluated the pipeline on several biomedical corpora and compared it with existing tools that support cell-type extraction. Across these evaluations, CellExLink showed clear accuracy gains in both detecting cell-type entities and assigning correct standard identifiers. Together, these gains make CellExLink a powerful tool for extracting reliable standardized cell-type information from large collections of papers, supporting literature curation, relation extraction, knowledge graph construction, and studies of cell-type-specific roles in diseases, drug responses, and biological pathways.

## Introduction

Cells are the fundamental units of biology, and many other biological entities are interpreted in relation to specific cell types [1, 2]. The importance of cell types is particularly evident in biological processes and disease states that are governed by interactions among diverse cellular and molecular components [3–7]. The computational task of identifying these cell types from published literature is important for advancing biomedical research and healthcare-related computational modeling. As an example, interactions among many cell types within the tumor microenvironment, including tumor cells, immune cells, and stromal cells, influence disease progression and treatment response [8–11]. These interactions are modeled using quantitative systems pharmacology (QSP) approaches to predict tumor progression and response to treatment [12–16]. The predictive value of such models depends on the accuracy and completeness of the underlying tumor microenvironment interaction network. Since these networks are often constructed from biomedical literature, reliable cell-type extraction is an important first step toward building more accurate interaction networks and supporting downstream computational modeling [12, 14, 17].

The task of extracting cell-type entities from literature has two linked parts: 1) named entity recognition (NER), which identifies the entity in text, such as cell types or cell populations; 2) named entity normalization (NEN), also called entity linking, which maps each entity to a standard identifier. The entity normalization step is needed because the same concept can appear in many different forms in text. For example, a cell type may be represented by several different names, such as (“granulosa cell”, “GC”), and (“endothelial cell”, “endotheliocyte”). Normalization maps these different forms to the same Cell Ontology (CL) identifier [18], which improves consistency across documents and supports downstream analysis.

Current off-the-shelf biomedical text-mining tools offer useful capabilities, but their support for cell-type recognition and normalization remains uneven. PubTator3 [19] recognizes genes/proteins, chemicals, diseases, species, variants, and cell lines, but not cell types. Another method, HunFlair2 [20], supports cell lines, chemicals, diseases, genes, and species but does not target cell types. GNorm2 [21], which is a strong system for gene recognition and normalization, is limited to genes. VANER2 [22] expands biomedical NER coverage, but it is a recognition system and does not link entities to ontology identifiers. Although a more comprehensive method, BERN2 [23], includes both cell-type recognition and normalization, its normalization strategy relies on rule-based methods, which have limited coverage. Another useful informatics software tool, ScispaCy [24], provides biomedical NER models with cell-related labels. However, its built-in linker does not support the Cell Ontology. As a result, an external integration such as PyOBO [25] is needed for CL linking. Here, we present an informatics tool that addresses an unmet need for practical, end-to-end support for both cell-type recognition and Cell Ontology normalization.

Furthermore, the advances in biomedical research indicate the need for new biomedical entity recognition tools. For instance, recent advances in single-cell technologies have changed how authors describe cell phenotypes [2, 26, 27]. Earlier corpora with cell-related annotations, including AnatEM [28], CRAFT [29], JNLPBA [30], and BioID [31], are still useful. However, recent work by Rotenberg et al. [27] suggests that these corpora do not capture the full range of fine-grained phenotypes reported in newer single-cell-focused papers. In that study, CellLink, built from literature published between 2019 and 2024, reported more than 22,000 annotated entities, 9,804 unique entities, and 1,251 unique Cell Ontology identifiers, covering nearly half of the current Cell Ontology. By comparison, BioID and CRAFT together contain 378 unique Cell Ontology identifiers. This shift suggests that systems trained only on earlier resources may not be able to identify many cell types found in recent papers and may not provide broad enough coverage of Cell Ontology identifiers.

The choice of model is an important consideration in developing an informatics tool for cell-type recognition and normalization. Recent advances in deep learning have made pretrained transformer models central to biomedical NLP. These models include encoder-based biomedical language models such as PubMedBERT and Bioformer, as well as generative large language models [32–37]. However, recent studies suggest that generative LLMs do not consistently outperform strong fine-tuned encoder-based biomedical models on information extraction tasks, although they can be competitive on reasoning-oriented tasks such as question answering [22, 38, 39]. In addition, they are generally more computationally expensive at inference [22]. Furthermore, NER and NEN are often the first stages of larger biomedical text-mining pipelines and are applied repeatedly across large corpora. Thus, their efficiency can affect the runtime of the overall pipeline [33]. For this reason, the design of cell-type extraction systems needs to consider computational efficiency alongside performance, motivating the use of fine-tuned encoder-based biomedical transformers.

Guided by these considerations, we developed CellExLink, an end-to-end pipeline for cell-type named entity recognition and Cell Ontology normalization. CellExLink adapts encoder-based biomedical language models by combining a jointly fine-tuned Bioformer [33] recognizer, a fine-tuned SapBERT [40] linker and a task-specific normalization process, all of which leverage recently developed cell-type-specific data resources [27]. Unlike general biomedical text-mining tools, CellExLink is tailored specifically to cell-type extraction and is intended to provide accurate and reliable support for recognizing cell-type entities and linking them to standardized Cell Ontology identifiers. We also considered computational efficiency when selecting the final recognition and normalization backbones to support integration into larger literature-mining workflows.

## Methods

### Overview

CellExLink has two components: 1) cell-type recognition and 2) Cell Ontology normalization. The recognition component identifies cell-type spans in the text, and the normalization component maps each recognized entity to a Cell Ontology (CL) identifier (Fig. 1 summarizes the workflow).

**Fig 1.**
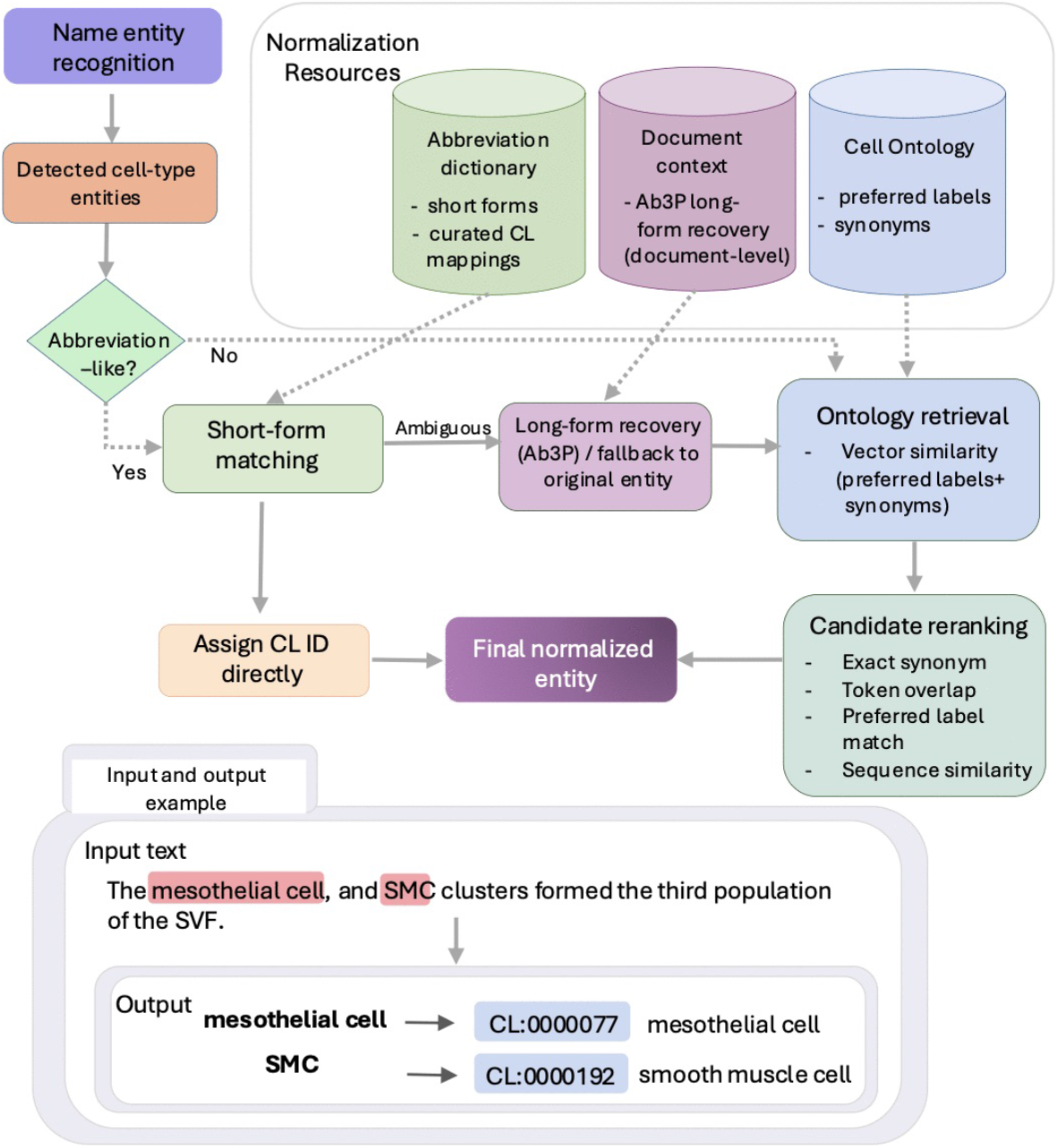
Overview of CellExLink. The pipeline detects cell-type entities in text and then links each entity to a Cell Ontology (CL) identifier.

### Cell-type recognition

To develop an accurate and efficient NER component, we evaluated two biomedical transformer backbones (PubMedBERT [32] and Bioformer16L [33]) under four fine-tuning strategies with different data compositions. The fine-tuning data included the CellLink [27] corpus and four earlier corpora: CRAFT [29], BioID [31], AnatEM [28], and JNLPBA [30], as summarized in Table 1.

**Table 1.** Datasets used in this study. Normalization and end-to-end evaluation were limited to datasets with CL identifiers. For CellLink, the validation set was used instead of the test set.

| Dataset | Text type | Origin / release | Corpus statistics (train / test) |  |  | CL ID |
| --- | --- | --- | --- | --- | --- | --- |
|  |  |  | Annotations | Unique entities | Documents |  |
| **CellLink** | Full-text article excerpts | Biomedical articles, 2019–2024 | 11,140 / 3,591 | 5,410 / 2,203 | 1,502 / 501 | Yes |
| **BioID** | Figure panel captions | SourceData / BioCreative VI Bio-ID, 2017 | 3,522 / 1,094 | 514 / 183 | 10,889 / 2,808 | Yes |
| **CRAFT** | Full-length journal articles | PubMed Central, 2012 release | 3,462 / 1,749 | 661 / 393 | 60 / 30 | Yes |
| **AnatEM** | Abstracts and text extracts | Extended anatomical entity corpus, 2014 | 1,787 / 1,031 | 766 / 530 | 603 / 202 | No |
| **JNLPBA** | MEDLINE abstracts | GENIA v3.02 shared-task corpus, 2004 | 6,710 / 1,909 | 2,141 / 940 | 1,999 / 403 | No |

To leverage the new CellLink dataset, we explored fine-tuning under different data compositions. Because CellLink better reflects recent literature on cell phenotypes than the earlier datasets, we treated it as a separate corpus, while merging the four earlier corpora into a single dataset representing prior training resources. Using this setup, we evaluated the following four fine-tuning strategies: (i) fine-tuning on CellLink alone; (ii) fine-tuning on the merged earlier corpora alone; (iii) fine-tuning on the merged earlier corpora, followed by further fine-tuning on CellLink; and (iv) joint fine-tuning on all corpora.

We selected the final recognition model based on its performance across five heterogeneous corpora. All corpora except CellLink include annotations for the test set. Therefore, for testing on CellLink data, we used its validation set. We computed corpus-level micro-F1 for both exact-span and relaxed-span matching and then macro-averaged these scores across corpora. We also considered computational efficiency in our analyses of these models.

### Cell-type normalization

We normalized each recognized entity to a Cell Ontology (CL) identifier using a task-specific method that combined dense retrieval, abbreviation handling, and candidate concept re-ranking. We used three normalization resources: (i) a CL alias inventory from BioPortal [41] based on the CL release dated 2025-12-17, (ii) an abbreviation dictionary derived from abbreviation-like entities in the fine-tuning data, and (iii) the input-document text available at inference time for document-level long-form recovery as shown in Fig. 1.

After evaluating multiple candidate encoders, we selected SapBERT [40] as the embedding model for NEN. We fine-tuned SapBERT on concept-labeled name sets built from CL aliases and CellLink [27] training entities mapped to CL identifiers. At inference time, we retrieved candidate aliases by comparing the entity embedding with alias embeddings using cosine similarity. We then grouped the retrieved aliases by concept and kept the highest-scoring alias for each concept.

For abbreviation-like entities, we first queried the abbreviation dictionary. We assigned unambiguous matches directly to their corresponding CL identifiers. If a match was ambiguous, we used Ab3P [42] to recover a long-form expansion from the input document text. We then passed the expanded form through the same candidate retrieval process. If no reliable expansion was available, we used the original entity for normalization.

After retrieving candidate CL concepts, we reranked them using the vector similarity score together with lexical features, including exact synonym match, token overlap, preferred-label overlap, and sequence similarity. We then assigned the top-ranked concept as the final normalized output.

## Results and discussion

We evaluated CellExLink at three levels: (i) cell-type recognition, (ii) Cell Ontology normalization on gold-standard entity spans, and (iii) strict end-to-end extraction, which required both a correct entity span and a correct ontology identifier. We first present the recognition results and the experiments used to select the final recognition backbone based on accuracy and efficiency. We then present the normalization results and the experiments used to select the final linker based on the same criteria. Finally, we present the strict end-to-end extraction results together with a case study.

### Cell-type recognition

Table 2 presents cell-type recognition performance across the five corpora. Table 3 summarizes the off-the-shelf tools that support cell-type recognition. We evaluated performance under both exact-span and relaxed-span criteria. Exact-span evaluation required an exact boundary match between the predicted and gold entity spans. Relaxed-span evaluation counted a prediction as correct when there was an overlap with the gold span at the token level. For example, a prediction of “T cells” was counted as correct for the gold span “CD8+ T cells”. To avoid giving credit to overly generic partial predictions under the relaxed matching, we blacklisted generic terms such as *cell* and *cellular*. For example, the prediction “cells” was not counted as correct for the gold entity “PIN cells”.

**Table 2.** Cell-type recognition performance on five corpora, measured by micro-F1 under exact-span (E) and relaxed-span (R) matching. Macro average denotes the average of per-corpus micro-F1 across the five corpora.

| System | CellLink |  | CRAFT |  | BioID |  | AnatEM |  | JNLPBA |  | Macro avg. |  |
| --- | --- | --- | --- | --- | --- | --- | --- | --- | --- | --- | --- | --- |
|  | E | R | E | R | E | R | E | R | E | R | E | R |
| ScispaCy | 0.308 | 0.488 | 0.375 | 0.462 | 0.136 | 0.181 | 0.248 | 0.334 | 0.222 | 0.382 | 0.258 | 0.369 |
| BERN2 | 0.684 | 0.847 | 0.406 | 0.592 | 0.244 | 0.413 | 0.567 | 0.731 | 0.663 | 0.778 | 0.513 | 0.672 |
| VANER2 | 0.430 | 0.837 | 0.180 | 0.566 | 0.315 | 0.452 | 0.302 | 0.813 | 0.318 | 0.741 | 0.309 | 0.682 |
| **CellExLink** | **0.860** | **0.941** | **0.740** | **0.840** | **0.745** | **0.800** | **0.764** | **0.864** | **0.723** | **0.832** | **0.766** | **0.855** |

**Table 3.** Representative tools for literature-based cell-type extraction compared in this study. Check marks indicate supported capabilities.

| System | NER | NEN | Backbone (params) | Year |
| --- | --- | --- | --- | --- |
| BERN2 | ✓ | ✓ | BioLM (~ 365M) | 2022 |
| VANER2 | ✓ | × | Llama 3.1 Instruct (~ 8B) | 2026 |
| ScispaCy | ✓ | ✓ | SpaCy pipeline | 2019/2025(updated) |
| **CellExLink** | ✓ | ✓ | BioFormer + SapBERT (~ 150M) | 2026 |

Across the five evaluation corpora, CellExLink outperformed all off-the-shelf baselines under both exact-span and relaxed-span evaluation (Table 2). BERN2 was the strongest off-the-shelf baseline overall under exact-span evaluation. VANER2 was more competitive under relaxed-span evaluation, which suggests that many of its errors were boundary errors rather than missed detections.

Fine-tuning data composition had a larger effect than encoder choice alone as shown in Fig 2. Models fine-tuned only on CellLink transferred less well to the earlier corpora. For both PubMedBERT and Bioformer16L, joint fine-tuning on all five corpora produced the best overall results (Fig. 2). The two jointly fine-tuned models performed similarly. However, Bioformer16L had a smaller model size and lower latency than PubMedBERT, as shown in Table 4. For this reason, we selected the jointly fine-tuned Bioformer16L model as the final NER model for CellExLink.

**Table 4.**
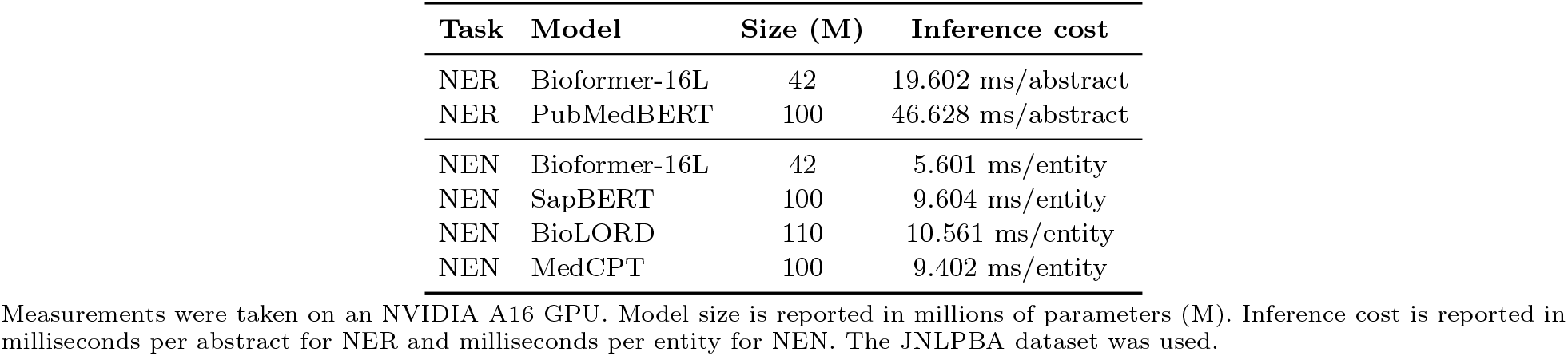
Model size and inference cost of candidate backbones.

| Task | Model | Size (M) | Inference cost |
| --- | --- | --- | --- |
| NER | Bioformer-16L | 42 | 19.602 ms/abstract |
| NER | PubMedBERT | 100 | 46.628 ms/abstract |
| NEN | Bioformer-16L | 42 | 5.601 ms/entity |
| NEN | SapBERT | 100 | 9.604 ms/entity |
| NEN | BioLORD | 110 | 10.561 ms/entity |
| NEN | MedCPT | 100 | 9.402 ms/entity |

Measurements were taken on an NVIDIA A16 GPU. Model size is reported in millions of parameters (M). Inference cost is reported in milliseconds per abstract for NER and milliseconds per entity for NEN. The JNLPBA dataset was used.

**Fig 2.**
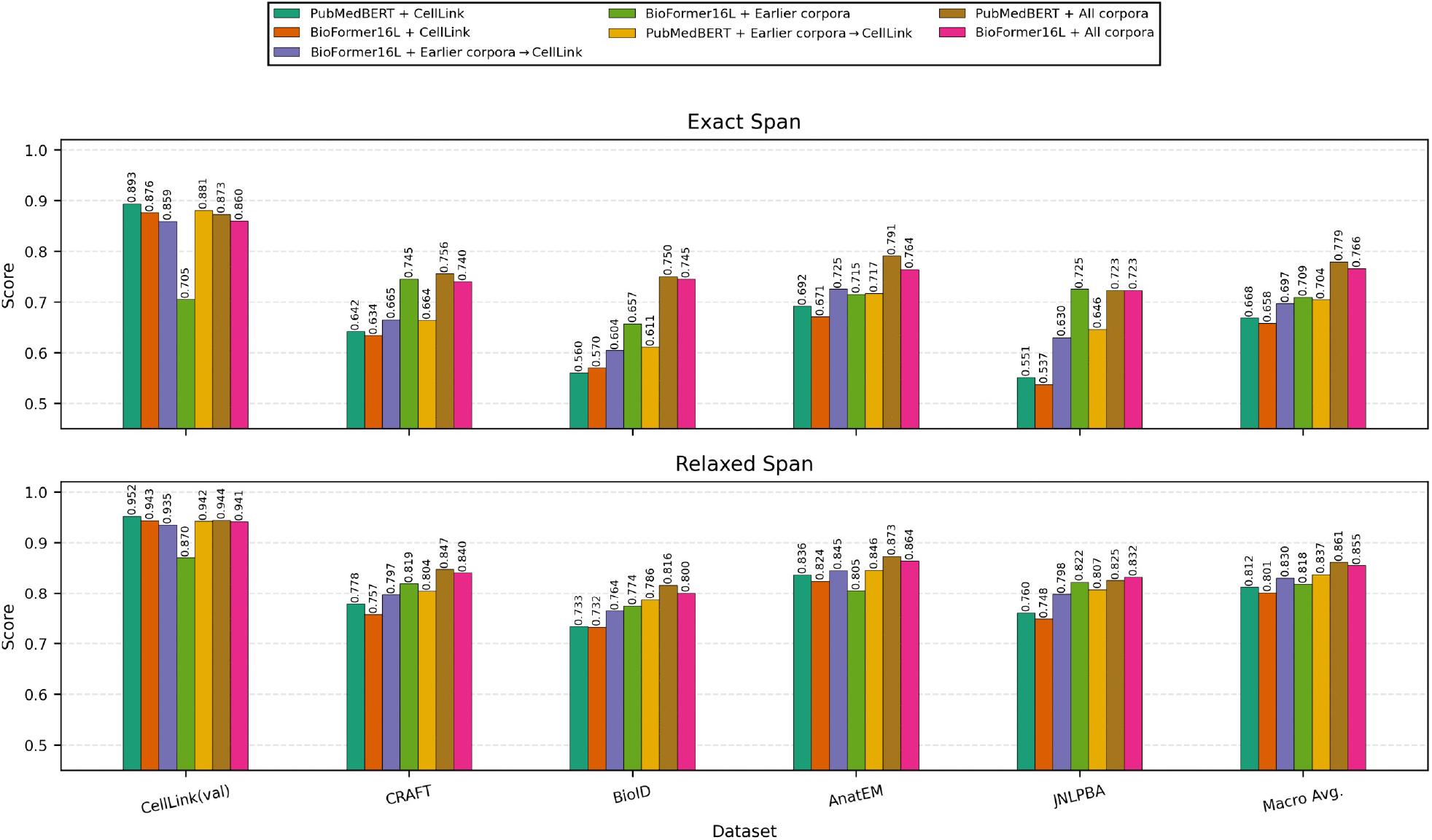
Effect of fine-tuning data and encoder choice on cell-type recognition. The top panel shows exact-span micro-F1, the bottom panel shows relaxed-span micro-F1, and the rightmost column in each panel shows macro-F1.

### Cell-type normalization on gold-standard spans

Table 5 presents the normalization results on gold-standard entity spans, which isolates linking performance from recognition errors. Under this setting, CellExLink outperformed both BERN2 and the ScispaCy baseline in every evaluation condition. The gains were largest on CellLink exact-match evaluation. CellExLink also remained better than both baselines on CRAFT and BioID.

**Table 5.** Cell-type normalization performance on gold-standard entity spans, reported as F1.

| Dataset | BERN2 | SciSpacy | CellExLink |
| --- | --- | --- | --- |
| **CellLink (exact matches only)** |  |  |  |
| Cell phenotype | 0.342 | 0.577 | **0.872** |
| Heterogeneous cell population | 0.356 | 0.544 | **0.939** |
| Overall | 0.342 | 0.576 | **0.874** |
| **CellLink (all IDs)** |  |  |  |
| Cell phenotype | 0.285 | 0.515 | **0.786** |
| Heterogeneous cell population | 0.174 | 0.363 | **0.601** |
| Overall | 0.278 | 0.506 | **0.775** |
| CRAFT | 0.331 | 0.490 | **0.818** |
| BioID | 0.220 | 0.457 | **0.690** |

Table 6 reports the performance of different linker backbones. The results show that no single zero-shot dense-retrieval backbone performed best across all datasets. SapBERT gave the strongest zero-shot result on CellLink exact matches and provided the best overall balance across datasets. MedCPT-Query [43] was slightly better on CRAFT, whereas BioLORD [44] performed best on BioID. Based on these results, we selected SapBERT as the starting point for further adaptation.

**Table 6.** Linker-selection results for cell-type normalization on gold-standard entity spans, reported as F1.

| Linker | CellLink exact | CellLink all | CRAFT | BioID |
| --- | --- | --- | --- | --- |
| *Zero-shot baselines* |  |  |  |  |
| SapBERT | 0.787 | 0.662 | 0.750 | 0.534 |
| MedCPT-Query | 0.750 | 0.653 | 0.755 | 0.571 |
| BioLORD | 0.752 | 0.664 | 0.713 | 0.647 |
| *Fine-tuned / adapted variants* |  |  |  |  |
| Bioformer16 + finetune | 0.783 | 0.692 | 0.743 | 0.620 |
| SapBERT + finetune | 0.825 | 0.728 | 0.802 | 0.645 |
| CellExLink | **0.874** | **0.775** | **0.818** | **0.690** |

We also evaluated a fine-tuned BioFormer16L as a linker backbone, because it was the most efficient retrieval backbone in our runtime comparison, as shown in Table 4. However, the fine-tuned SapBERT model gave better normalization accuracy overall; we therefore continued with SapBERT. As shown in Table 6, fine-tuning improved performance over the zero-shot SapBERT model. The full CellExLink normalization process, which used fine-tuned SapBERT together with the full inference procedure, produced further gains.

### Strict end-to-end extraction

Table 7 reports strict end-to-end results, i.e., a prediction was counted as correct only when both the entity span and the Cell Ontology identifier were correct. This setting provides a practical end-to-end evaluation for downstream literature mining. CellExLink substantially outperformed both baselines across all three evaluation datasets. The gap between gold-span normalization and end-to-end extraction indicates that recognition errors remained an important source of performance loss. Much of this loss came from missed mentions and span-boundary errors, which reduced recall. Overall, CellExLink still outperformed both baseline tools by a large margin.

**Table 7.**
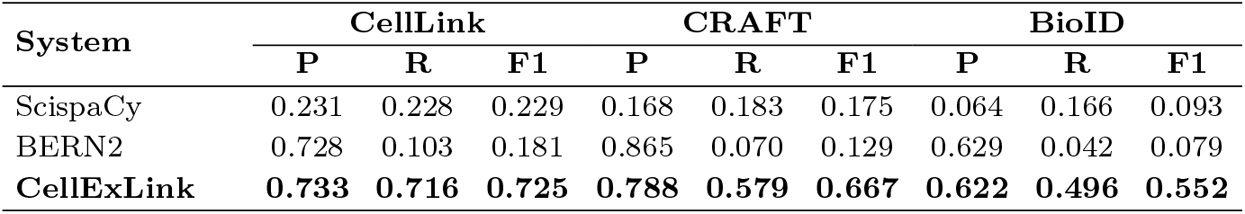
Strict end-to-end cell-type extraction performance. A prediction was counted as correct only when both the mention span and the Cell Ontology identifier were correct. Scores are reported as linked precision (P), recall (R), and F1.

| System | CellLink |  |  | CRAFT |  |  | BioID |  |  |
| --- | --- | --- | --- | --- | --- | --- | --- | --- | --- |
|  | P | R | F1 | P | R | F1 | P | R | F1 |
| ScispaCy | 0.231 | 0.228 | 0.229 | 0.168 | 0.183 | 0.175 | 0.064 | 0.166 | 0.093 |
| BERN2 | 0.728 | 0.103 | 0.181 | 0.865 | 0.070 | 0.129 | 0.629 | 0.042 | 0.079 |
| **CellExLink** | **0.733** | **0.716** | **0.725** | **0.788** | **0.579** | **0.667** | **0.622** | **0.496** | **0.552** |

### Case Study

We also conducted a case study comparing CellExLink with the off-the-shelf tools ScispaCy and BERN2. The examples are shown in Fig 3. In case 1, CellExLink correctly recovered the coordinated and modifier-rich entity “slow and fast muscle fibres”, whereas both other tools failed. In cases 2 and 3, CellExLink also correctly detected and linked “CD8+ T cells” and “trophoblast progenitor cells”. In contrast, ScispaCy linked these entities to broader concepts such as “T cells” and “trophoblast”, whereas BERN2 detected the spans but did not assign Cell Ontology identifiers. These examples show that the gains of CellExLink came from improvements in both span detection and ontology linking.

**Fig 3.**
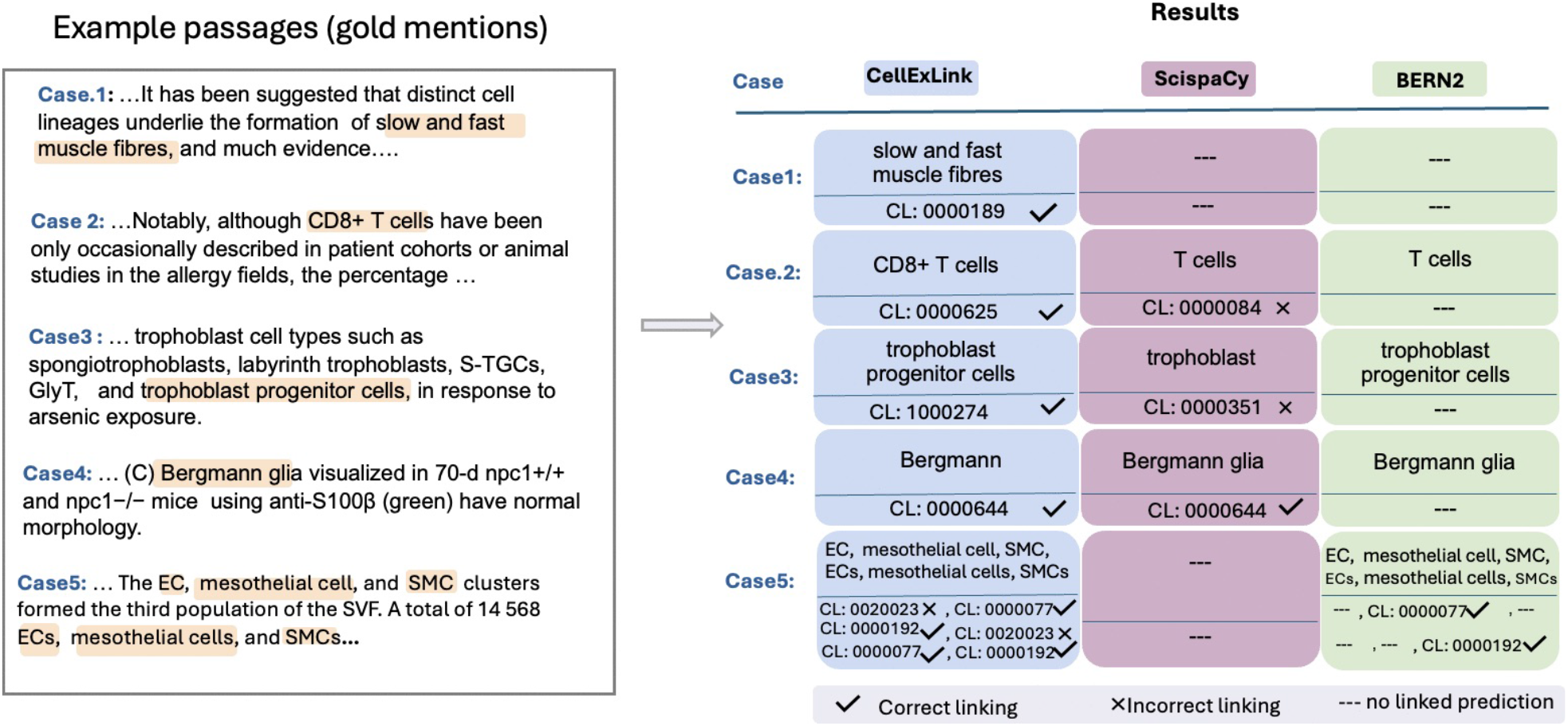
A case study comparing CellExLink with ScispaCy and BERN2. The left panel shows example passages and gold mentions. The right panel shows the mentions and Cell Ontology identifiers returned by each system.

There are still several limitations in our work, particularly for short figure captions, where context was sparse and abbreviations were frequent. For example, in case 5, CellExLink linked “mesothelial cell”/”mesothelial cells” and “SMC”/”SMCs” consistently, but linked “EC”/”ECs” to incorrect Cell Ontology identifiers. In another case, CellExLink split a gold entity into smaller spans within the caption text. These errors show that short abbreviations and exact boundary recovery remain difficult in a short, caption-style text with limited surrounding context. Overall, CellExLink achieved strong general performance across all datasets.

## Conclusion

In this work, we presented CellExLink, an end-to-end tool for literature-based cell-type extraction that integrates cell-type recognition with Cell Ontology normalization. Across multiple evaluation corpora, CellExLink improved cell-type recognition, normalization, and strict end-to-end extraction relative to currently available off-the-shelf tools that support cell-type extraction.

This work has some limitations that can be overcome in future studies. For example, the current annotation setup used flat, non-overlapping entity annotations, which may not fully represent overlapping, nested entities. Because of this, some entities may have been simplified or not fully captured. Short figure captions and abbreviation-heavy text also remained challenging, because they provide limited context for both span detection and normalization. In the future, the cell-type recognition and normalization methods can be improved by supporting richer annotation structures, enhanced abbreviation handling, and stronger boundary detection.

## 1 Data and code availability

Information for accessing CellExLink, including the source code, datasets, documentation, and model checkpoints, is available on GitHub at https://github.com/ShahriyariLab/CellExLink-End-to-End-Cell-Type-Extraction-and-Cell-Ontology-Normalization-from-Biomedical-Text

